# Amygdala habituation during exposure is associated with failure to reduce phobic symptoms

**DOI:** 10.1101/2023.12.29.573600

**Authors:** Cindy S. Lor, Alexander Karner, Mengfan Zhang, Kathrin Kostorz, Ronald Sladky, Frank Scharnowski

**Affiliations:** Department of Cognition, Emotion, and Methods in Psychology, University of Vienna, Liebiggasse 5, 1010 Vienna, Austria

**Author notes:** <u>Corresponding author:</u> Cindy S. Lor.

## Abstract

Exposure therapy is an established treatment for anxiety disorders but the mechanisms underpinning its effectiveness remain unclear. Two theories offer contrasting perspectives: the traditional habituation model posits that a form of stimulus desensitization is required during exposure, while the inhibitory learning model emphasizes the formation of new non-fearful associations. Crucially, while the former may manifest as amygdala habituation, the latter may not. To distinguish between the two models, this study uses functional magnetic resonance imaging to examine the amygdala responses of spider-fearful participants during fear exposure. We hypothesized that intervention success might align with stable or even increased amygdala activation – an indicator of active engagement and re-learning as proposed by the inhibitory learning model. Conversely, decreasing amygdala activity might not be a sign of reduced fear memory as proposed by the habituation model, but could signal mental detachment, leading to suboptimal treatment outcomes. Our results corroborated our hypotheses: individuals with escalating amygdala responses during exposure exhibited better clinical progress, while those showing amygdala ‘habituation’ benefited less. Our results strengthen the case for the inhibitory learning model and highlight that therapy may not aim to diminish fear per se but rather to engage patients in active processing and association formation.

## 1. Introduction

Exposure therapy is a well-established therapeutic approach for anxiety disorders that are linked to distinct and identifiable external triggers. During exposure therapy, individuals are methodically and progressively exposed to problematic stimuli, allowing them to develop strategies to mitigate associated fear and anxiety reactions (Andrews et al., 2002). Such therapy has proven especially effective for disorders like specific phobias, social anxiety disorder, and agoraphobia, wherein fear responses are triggered by particular situations, objects, or contexts. Specific phobias are one of the most common anxiety disorders globally and present a substantial challenge for many individuals (Wardenaar et al., 2017). Spider phobia is particularly widespread, with estimates ranging from 2.7 to 9.5% (Oosterink et al., 2009; Zsido, 2017).

Despite generally high treatment efficacy for specific phobias, not all individuals who undergo exposure therapy benefit from it and relapse rates are considerable (Böhnlein et al., 2020; Thng et al., 2020). Also, the efficacy of exposure therapy for other types of anxiety disorders and for more multifaceted disorders, such as post-traumatic stress disorders (PTSD), tends to be lower (Markowitz & Fanselow, 2020). These limitations raise questions about the therapy’s underlying mechanisms and its applicability spectrum. Studying the boundaries of its effectiveness using neuroscientific tools could provide insights into optimizing therapeutic strategies and improve the range of applications.

The amygdala is a central structure to the neurobiology of exposure therapy, and is well-known for its involvement in processing and regulating fear and anxiety (Garcia, 2017). Within the amygdala, the basolateral complex (BLA) and the central nucleus (CeA) are of particular interest due to their distinct, albeit interconnected, roles (Hartley et al., 2019). The BLA is involved in the cognitive evaluation of threats, integrates sensory information and communicates with other brain areas like the prefrontal cortex to assign emotional significance to stimuli. This evaluative process integrates prior experiences, context, and other cognitive factors to shape the emotional response. Additionally, the BLA plays a pivotal role in fear conditioning, where it is involved in forming associations between neutral stimuli and aversive events (Fanselow & LeDoux, 1999; Hartley et al., 2019). The CeA may be more involved in orchestrating autonomic and behavioral responses to fear (Keifer et al., 2015). Upon receiving information from the BLA, the CeA is thought to activate various physiological responses such as increased heart rate, blood pressure, and stress hormone release (Tsigos et al., 2000).

Several neuroscientific studies have associated a reduction of amygdala activity with successful exposure therapy (Goossens et al., 2007; Hauner et al., 2012). However, the majority of these investigations have focused on the contrast between pre- and post-treatment effects instead of the dynamics within the exposure session. A notable exception studied brain activity during exposure, and found that the decreased activation in one cluster in the amygdala correlated with positive clinical outcome in spider-phobic individuals (Björkstrand et al., 2020). This finding is in line with the habituation model which posits that, after fear activation, repeated exposure to a feared stimulus would lead to a decline in phobia symptoms, and this decline is seen as an essential factor of the therapeutic process (Benito & Walther, 2015).

Yet, recent years have seen a paradigm shift, with a growing emphasis on the inhibitory learning model (Craske et al., 2014; Tolin, 2019; Weisman & Rodebaugh, 2018). This model proposes that the success of exposure therapy is rooted in building new, non-fearful associations that may coexist with the previous fearful ones. Contrary to the habituation model, this model proposes that the goal of exposure therapy is not necessarily to reduce fear during the exposure but to facilitate the learning of new, safety associations that can compete with older, fearful ones. Intriguingly, while the habituation model might interpret reduced amygdala activity as a sign of therapeutic success, under the inhibitory learning model, one could perceive it as a sign of dissociation or reduced cognitive engagement, potentially undermining therapy’s effectiveness.

In this study, we first aimed to investigate the behavior of the whole amygdala during the exposure session. We hypothesized that a sustained, or even heightened, activity could indicate effective therapeutic engagement and be linked to clinical improvement. As a second objective, we turned our focus to the amygdala’s sub-divisions, as they may capture distinct functions and may show different dynamics. Since the BLA seems to be more closely related to the formation of novel fear associations, we expect it to mirror the behavior of whole amygdala. On the other hand, we hypothesized that the activity of the CeA, a much smaller structure related to autonomic responses, may decrease within the session as a sign of improved phobia symptoms. This would be in line with the habituation model, suggesting an alleviation of phobia symptoms.

To validate these hypotheses, we analyzed functional magnetic resonance imaging (fMRI) data of a population of sub-clinical spider-fearful individuals while they were exposed to pictures of spiders. We also collected pre-post exposure phobia severity scores (Fear of Spider Questionnaire (FSQ) (Szymanski & O’Donohue, 1995) and Behavioral Avoidance Test (BAT) (Muris et al., 1998)) one day (up to one week) after scanning (follow-up-1) and one month after scanning (follow-up-2). We extracted neural activity from the whole amygdala, the BLA and the CeA and correlated the change of activity over the course of the session with their change of clinical behavior. We expect clinical improvement to be positively associated with increased BLA/whole amygdala activity, while its association with the change of CeA should be negative.

## 2. Methods

### 2.1. Participants

Forty-nine German-speaking individuals (age = 22.6±3.5, 40F, 9M) with at least moderate spider fear levels (a minimum of 8 out of 24 points on the German Spider Fear Screening Questionnaire (SAS, (Rinck et al., 2003); SAS = 16.1±4.4) participated in the study. Exclusion criteria included psychiatric history, pregnancy, and substance use. Participants received a total compensation of 50€, with an added 5€ COVID-testing compensation fee when needed. Data was collected between September 2021 and November 2022. The study adhered to the Declaration of Helsinki and received approval from the University of Vienna’s ethics committee (IRB numbers: 00584, 00657).

### 2.2. fMRI paradigm

Participants underwent a brain scan at the University of Vienna’s MR Center. First, a resting-state scan was performed during which participants were instructed to relax and keep their eyes open and directed at a fixation point. Subsequently, participants watched a set of 225 spider and 45 non-spider neutral pictures divided into five runs. Each image presentation lasted for 4 seconds and was followed by a presentation of a 2-3 sec jittered fixation point. Six button-press catch trials per run were also included to ensure participants’ engagement. Between runs, we asked participants about their current levels of agitation and exhaustion, which they stated via the intercom on a scale from 0 to 10 to further maintain their engagement. The five passive-viewing runs were followed by another resting-state scan. An anatomical scan was acquired at the end of the scanning session. To reduce head motion, we used tape which provided tactile feedback for motion on the forehead as described by (Krause et al., 2019).

### 2.3. Pre-post behavioral assessment

We used the BAT and the FSQ to assess levels of spider fear severity. The FSQ was completed three times (1) a few days prior to the scanning session, online (pre-exposure), (2) a few days after the scanning session, online (follow-up-1) and (3) about a month after the scanning session, in person (follow-up-2).

The BAT was completed twice, once just before scanning (pre-BAT) and a second time after scanning at follow-up-2. Participants were instructed to approach a terrarium containing an inanimate but alive-looking huntsman spider until they still felt comfortable, and may open the lid of the terrarium if they dared to. BAT scores were measured as the total distance travelled to get closer to the spider, with lower BAT values indicating more fear. In contrast, lower FSQ scores indicate less fear. After the first BAT (pre-exposure), the participants walked for about 15 minutes to the scanning site to allow for wash out effects. A second BAT was performed about a month after the scanning session (post-exposure follow-up-2) (see Figure 1 for design). For a small subset of the participants (9/49) the sequence of the FSQ and BAT was reversed. Although this may have introduced priming effects that could have confounded the FSQ outcomes, we observed no statistical difference in any clinical measure between this subgroup and the rest of the sample.

**Figure 1.**
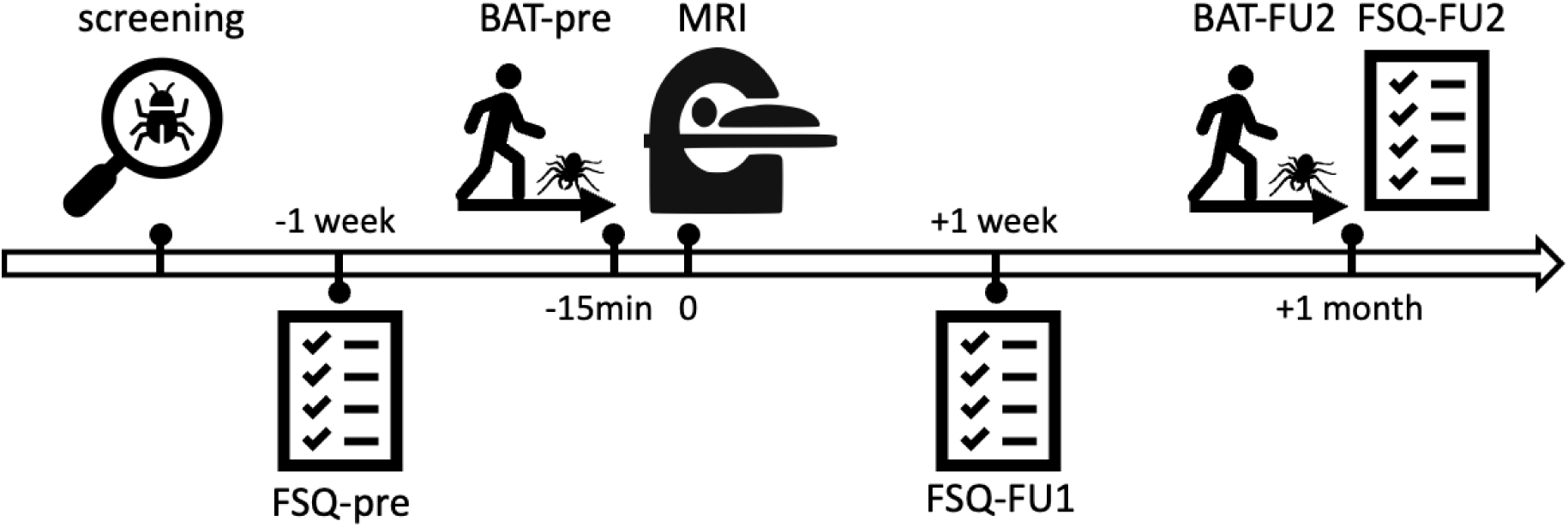
Timeline of the experimental procedure. Following inclusion after screening, the participants completed the FSQ questionnaire approximately one week before the scanning session. Immediately prior to the scanning session, the participants performed a BAT. At follow-up-1 (FU1), one week after scanning, participants completed another FSQ questionnaire, and at follow-up-2 (FU2), approximately 1 month later, they completed another BAT and FSQ.

While the study did not have explicit clinical trial-like objectives with strong expectations for clinical improvement, we employed one-sided paired t-tests to evaluate changes in FSQ and BAT scores at the group level.

### 2.4. MRI Acquisition

Data was acquired using a 3T Siemens MAGNETOM Skyra MRI scanner (Siemens, Erlangen, Germany) with a 32-channel head coil, at the University of Vienna. Functional data were collected with an interleaved mode, using a multiband factor 4 accelerated T2*-weighted echo planar imaging sequence, TR = 1250 ms, TE = 36 ms, voxel size = 2 mm x 2 mm x 2.6 mm, flip angle = 65 degrees, field of view (FOV) = 192 mm x 192 mm x 145 mm, 56 slices without slice gap to cover the full brain, anterior-posterior phase encoding. The total scan duration for the resting states (not analyzed in this study) was 8:51 min, and 7:11 min for the passive viewing per run (345 TRs per run). Structural data for each participant was acquired using a single-shot, high-resolution MPRAGE sequence with an acceleration factor of 2 using GRAPPA: TR = 2300 ms. TE = 2.43 ms, voxel size = 0.8 mm x 0.8 mm x 0.8 mm, flip angle = 8 degrees, field of view (FOV) = 263 mm x 350 mm x 350 mm.

### 2.5. MRI Preprocessing

Pre-processing was performed using fMRIPrep 20.2.6 (Esteban et al., 2019), which is based on Nipype 1.7.0 (Gorgolewski et al., 2011).

#### 2.5.1 Anatomical data preprocessing

First, T1-weighted (T1w) images were corrected for intensity non-uniformity (INU) and defined as T1w-reference for the following workflow. After skull-stripping with antsBrainExtraction, brain images were segmented into cerebrospinal fluid (CSF), white-matter (WM) and gray-matter (GM) using Fast (Zhang et al., 2001). Volume-based spatial normalization to the standard space (MNI152NLin2009cAsym) was performed through nonlinear registration using antsRegistration (ANTs 2.3.3), with brain-extracted versions of both the T1w reference and T1w template.

#### 2.5.2. Functional data preprocessing

The functional preprocessing with fMRIPrep included creating a reference volume, followed by skull-stripping. The reference then underwent distortion correction utilizing an estimated phase-difference fieldmap. The reference volume was aligned to the participant’s T1w space using FSL tools (version 5.0.9). This alignment was performed with nine degrees of freedom. Subsequently, the head-motion parameters, with respect to the reference volume, were calculated using MCFLIRT in FSL (version 5.0.9), post slice-time adjustment of the BOLD sequences with the aid of AFNI. The adjusted BOLD sequences were then transformed to MNI space by combining the transformation parameters estimated earlier, using antsApplyTransforms from ANTs (version 2.3.3) set with Lanczos interpolation. aCompCor parameters were also computed from the WM and CSF masks. Spatial smoothing with a 6 mm FWHM Gaussian kernel was applied as the last pre-processing step using SPM12. Two runs with excessive head movement (max framewise displacement (FD) > 5) were excluded and were treated as missing data points.

### 2.6. First-level analysis

To extract brain activity, we followed a GLM-based procedure using SPM12 which ran on MATLAB_R2023a. We performed first-level analysis to obtain beta/activation maps, specifying three regressors, one for fear-inducing spider image trials, one for neutral image trials and one for button-press catch trials. Each regressor modelled the onset and duration of stimulus presentation as a boxcar convolved with the hemodynamic response function. The following nuisance regressors were also added: six head motion realignment parameters, the first five aCompCor components of the CSF as well as of the WM. Of note, our experimental design included spider pictures that weren’t anticipated to trigger significant fear reactions. These images were part of a broader experiment intended to focus on capturing a spectrum of fear responses, ranging from mild discomfort to intense phobia. For the current analysis, we concentrated on images that elicit potent fear reactions, i.e. they had to be rated above 27/100 on our empirical fear scale and ignored the ones below this threshold. This cut-off value was chosen prior to any analysis and ensured that we did not misattribute inanimate spider representations, like spider toys or cartoons, to either the “neutral” or “fear-inducing spider” categories.

### 2.7. Second-level analysis

Next, for each run, we computed spider > neutral contrast maps to capture the brain activity associated with viewing fear-inducing spiders as compared to non-spider neutral images. We generated contrast maps per run (one for each run). We then used custom Matlab scripts to extract the average contrast within an anatomical mask of the whole amygdala, the BLA and of the CeA (Tyszka & Pauli, 2016), to obtain ROI-averaged activities per run.

For quality assessment purposes of the analysis pipeline, we additionally generated contrast maps that average all five runs to perform a voxel-wise validation analysis on the whole brain. Here, we applied a one-sample group analysis of subject-wise contrast maps in SPM12, with a height threshold of p = 0.001, and a family-wise error-corrected cluster threshold of p < 0.05. The purpose of this analysis is to identify brain regions that are more activated during spider viewing as opposed to neutral picture viewing, which should reveal conventional “fear brain centers”. We used the AAL toolbox for cluster labelling (Rolls et al., 2020).

### 2.8. Amygdala x clinical improvement correlation analyses

We quantified the change of activity within the exposure session as the difference between the mean contrast of the two first runs and the two last runs. Short-term FSQ improvement was defined as the difference in FSQ between pre-exposure and follow-up 1 (pre minus follow-up-1). Long-term FSQ improvement was defined as the difference in FSQ between pre-exposure and follow-up 2 (pre minus follow-up-2). BAT improvement was defined as the difference in BAT scores between pre-exposure to follow-up 2 (follow-up-2 minus pre). To account for the fact that lower BAT scores represent more fear severity, while lower FSQ scores indicate less fear severity, we inverted the sign for the FSQ difference. Thus, in all cases, higher scores indicate higher improvement. The relationship between amygdala activity change and clinical improvement was tested using Spearman correlation. Some clinical data are missing mostly due to COVID19-related reasons or technical failure. Four participants did not complete the BAT either at pre-exposure BAT (one participant) or at follow-up-2 (three other participants). FSQ data is missing at FSQ follow-up-1 (ten participants), as well as FSQ follow-up-2 (three participants). In total, our sample sizes for clinical evaluation are N = 45 for BAT improvement, N = 39 for short-term FSQ improvement, and N = 46 for long-term FSQ improvement. For each ROI, to control for family-wise error rate, we applied the Bonferroni-Holm correction, which is uniformly more powerful than the Bonferroni correction.

### 2.9. Exploratory atlas-based correlation analyses

While our primary focus was on the amygdala’s role in exposure, to provide a comprehensive context and underscore the significance of our results, we applied our ROI-averaged analysis to all the regions of the AAL3 atlas, computed the difference between the two first and two last runs, and correlated this difference with the three measures of clinical improvement. This “control analysis” aimed to benchmark the correlation strength of the amygdala against other brain regions.

## 3. Results

### 3.1. Comparison of clinical behavior

We observed a significant improvement of BAT at follow-up-2 (mean = 3.14 meters ± 0.61) compared to BAT at pre-exposure (mean = 2.90 meters ± 0.73) (p = 0.0152). For FSQ, compared to pre-exposure (mean = 55.27± 20.93), there was no significant improvement at follow-up-1 (mean = 50.87 ± 19.10) (p = 0.1282), but the improvement was significant at follow-up-2 (mean = 47.34 ± 19.05) (p = 0.0013) (Figure 2).

**Figure 2.**
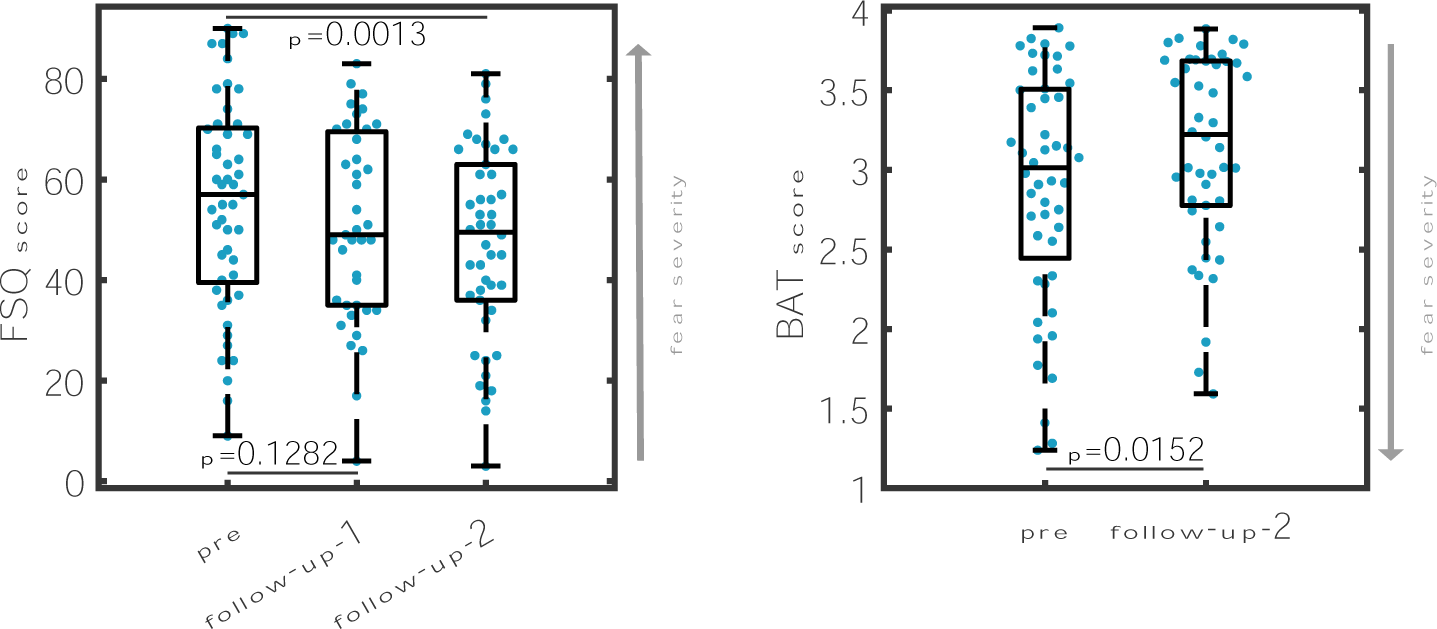
Clinical score comparison before and after the experimental exposure session (pre: before exposure; follow-up-1: ∼ a few days after the session; follow-up-2: ∼ a month after the session). FSQ = fear of spider questionnaire; BAT = behavioral avoidance test.

### 3.2. Whole-brain validation analysis

Our analysis revealed clusters of activation in most fear brain centers, including the bilateral amygdala, periaqueductal grey, hippocampus, putamen, insula, ACC, thalamus, cerebellum (Figure 3). See Table 1.

**Figure 3.**
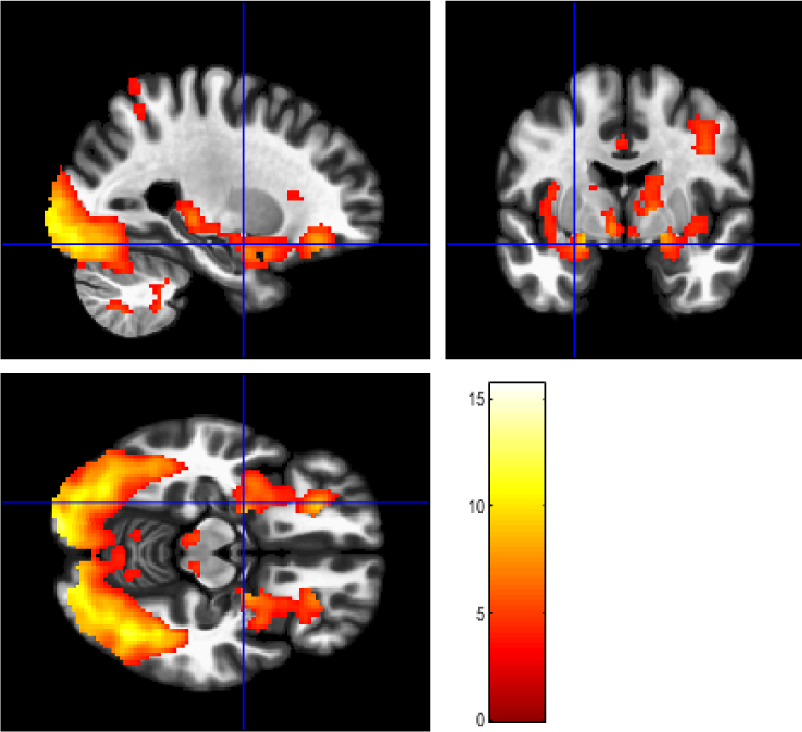
Whole brain map showing spider>neutral images activation clusters obtained from second-level one-sample t-test analysis (peak uncorrected threshold p = 0.001; FWE-corrected cluster threshold p = 0.05). N=49.

**Table 1.**
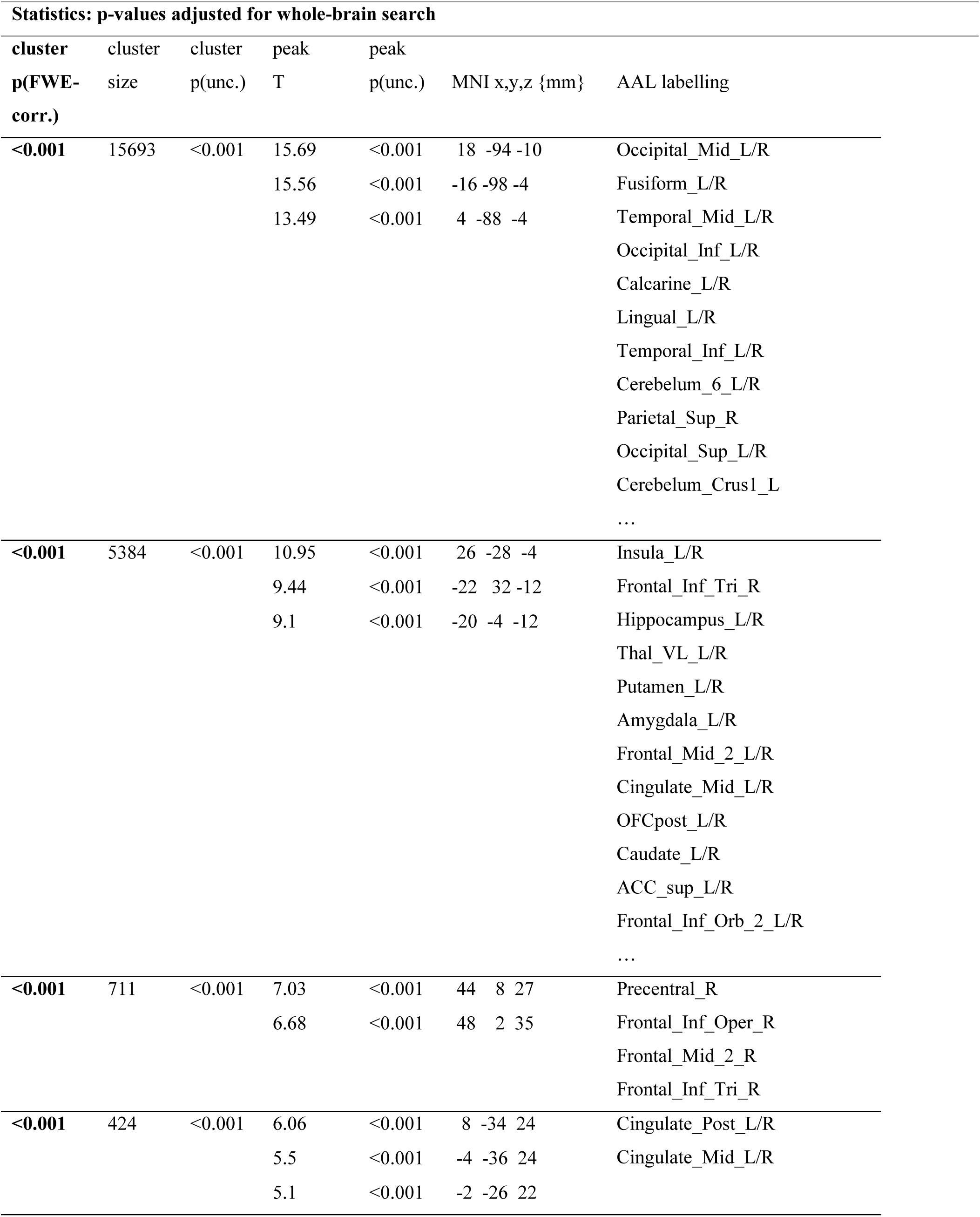
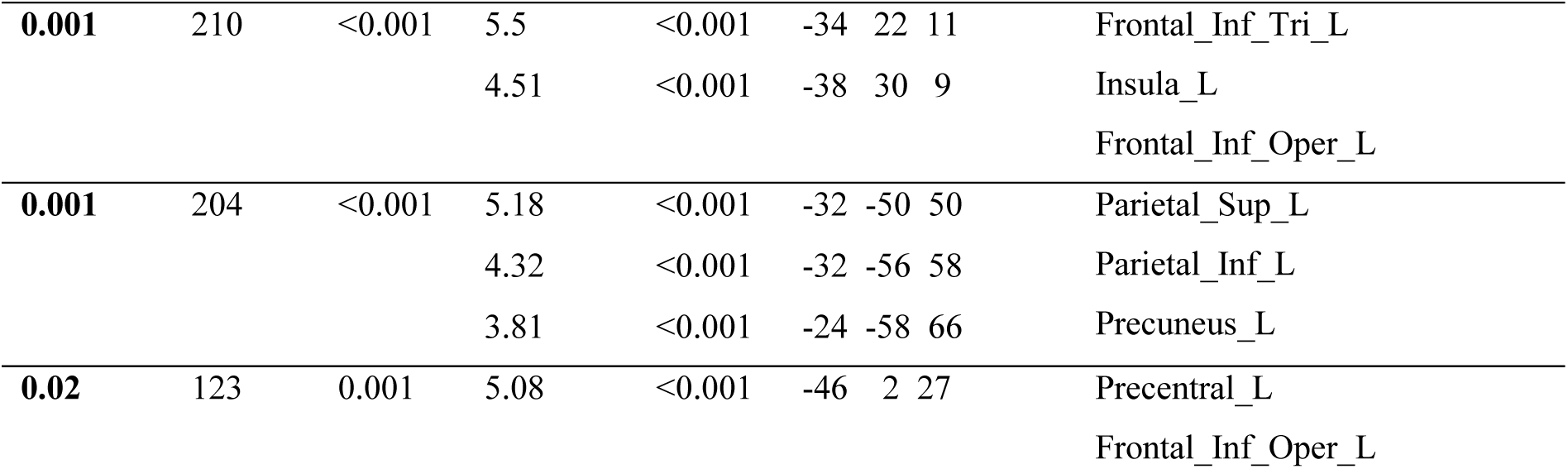
Whole-brain second-level results for spider>neutral contrast (peak uncorrected threshold p = 0.001; FWE-corrected cluster threshold p = 0.05). N=49.

| Statistics: p-values adjusted for whole-brain search |  |  |  |  |  |  |
| --- | --- | --- | --- | --- | --- | --- |
| cluster<br>p(FWE-corr.) | cluster<br>size | cluster<br>p(unc.) | peak<br>T | peak<br>p(unc.) | MNI x,y,z {mm} | AAL labelling |
| <0.001 | 15693 | <0.001 | 15.69 | <0.001 | 18 -94 -10 | Occipital_Mid_L/R |
|  |  |  | 15.56 | <0.001 | -16 -98 -4 | Fusiform_L/R |
|  |  |  | 13.49 | <0.001 | 4 -88 -4 | Temporal_Mid_L/R |
|  |  |  |  |  |  | Occipital_Inf_L/R |
|  |  |  |  |  |  | Calcarine_L/R |
|  |  |  |  |  |  | Lingual_L/R |
|  |  |  |  |  |  | Temporal_Inf_L/R |
|  |  |  |  |  |  | Cerebelum_6_L/R |
|  |  |  |  |  |  | Parietal_Sup_R |
|  |  |  |  |  |  | Occipital_Sup_L/R |
|  |  |  |  |  |  | Cerebelum_Crus1_L |
|  |  |  |  |  |  | ... |
| <0.001 | 5384 | <0.001 | 10.95 | <0.001 | 26 -28 -4 | Insula_L/R |
|  |  |  | 9.44 | <0.001 | -22 32 -12 | Frontal_Inf_Tri_R |
|  |  |  | 9.1 | <0.001 | -20 -4 -12 | Hippocampus_L/R |
|  |  |  |  |  |  | Thal_VL_L/R |
|  |  |  |  |  |  | Putamen_L/R |
|  |  |  |  |  |  | Amygdala_L/R |
|  |  |  |  |  |  | Frontal_Mid_2_L/R |
|  |  |  |  |  |  | Cingulate_Mid_L/R |
|  |  |  |  |  |  | OFCpost_L/R |
|  |  |  |  |  |  | Caudate_L/R |
|  |  |  |  |  |  | ACC_sup_L/R |
|  |  |  |  |  |  | Frontal_Inf_Orb_2_L/R |
|  |  |  |  |  |  | ... |
| <0.001 | 711 | <0.001 | 7.03 | <0.001 | 44 8 27 | Precentral_R |
|  |  |  | 6.68 | <0.001 | 48 2 35 | Frontal_Inf_Oper_R |
|  |  |  |  |  |  | Frontal_Mid_2_R |
|  |  |  |  |  |  | Frontal_Inf_Tri_R |
| <0.001 | 424 | <0.001 | 6.06 | <0.001 | 8 -34 24 | Cingulate_Post_L/R |
|  |  |  | 5.5 | <0.001 | -4 -36 24 | Cingulate_Mid_L/R |
|  |  |  | 5.1 | <0.001 | -2 -26 22 |  |

|  |  |  |  |  |  |  |  |  |
| --- | --- | --- | --- | --- | --- | --- | --- | --- |
| **0.001** | 210 | <0.001 | 5.5 | <0.001 | -34 | 22 | 11 | Frontal_Inf_Tri_L |
|  |  |  | 4.51 | <0.001 | -38 | 30 | 9 | Insula_L |
|  |  |  |  |  |  |  |  | Frontal_Inf_Oper_L |
| **0.001** | 204 | <0.001 | 5.18 | <0.001 | -32 | -50 | 50 | Parietal_Sup_L |
|  |  |  | 4.32 | <0.001 | -32 | -56 | 58 | Parietal_Inf_L |
|  |  |  | 3.81 | <0.001 | -24 | -58 | 66 | Precuneus_L |
| **0.02** | 123 | 0.001 | 5.08 | <0.001 | -46 | 2 | 27 | Precentral_L |
|  |  |  |  |  |  |  |  | Frontal_Inf_Oper_L |

### 3.3. Whole amygdala x clinical improvement correlation analyses

For the whole amygdala, we found a significant correlation between the change of amygdala contrast activity and all our measures of clinical improvement. One subject had to be excluded due to excess motion in the first 2 runs. For the whole amygdala, we found a significant correlation between the difference and all measures of clinical improvement (whole amygdala x short-term FSQ: r = 0.3749; p_unc_ = 0.0204; p_corr_ = 0.0204 - whole amygdala x long-term FSQ: r = 0.3768; p_unc_ = 0.0107; p_corr_ = 0.0214 - whole amygdala x BAT: r = 0.4584; p_unc_ = 0.0019; p_corr_ = 0.0057) (Figure 4).

**Figure 4.**
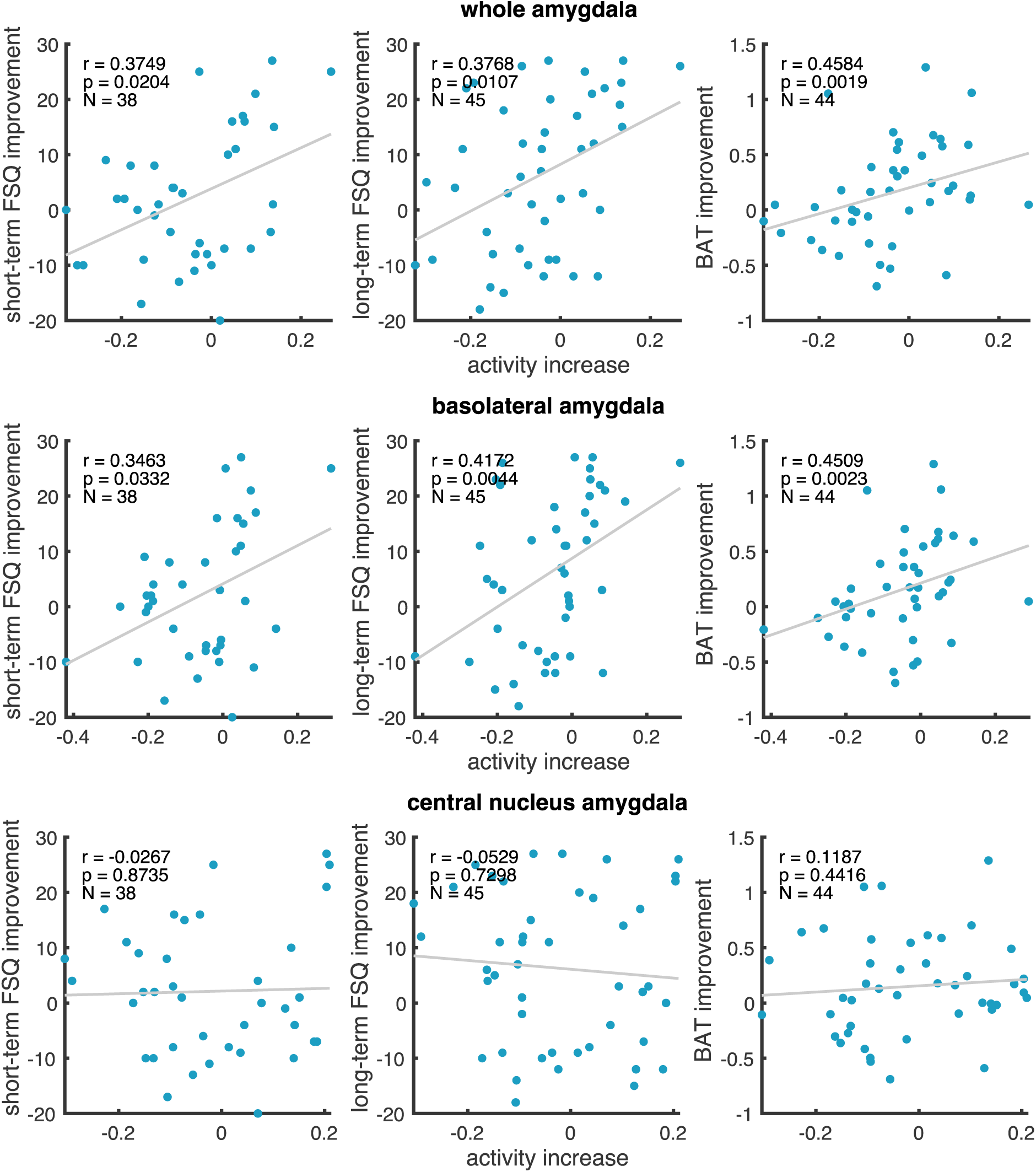
Correlation scatter plots between change in spider>neutral amygdala contrast (difference between the first and last two runs) and clinical improvement. Graphs show uncorrected p-values.

### 3.4. Amygdala subdivisions x clinical improvement correlation analyses

We observed similar results to the whole amygdala for the BLA (BLA x short-term FSQ: r = 0.3463; p_unc_ = 0.0332; p_corr_ = 0.0332 - BLA x long-term FSQ: r = 0.4271; p_unc_ = 0.0044; p_corr_ = 0.0088 - BLA x BAT: r = 0.4509; p_unc_ = 0.0023; p_corr_ = 0.0069). Finally, there was no correlation between the change in CeA contrast activity and neither of the three clinical measures (CeA x short-term FSQ: r = −0.0267; p_unc_ = 0.8735; p_corr_ = 0.8735 - CeA x long-term FSQ: r = −0.0529; p_unc_ = 0.7298; p_corr_ = 1 - CeA x BAT: r = 0.1187; p_unc_ = 0.4416; p_corr_ = 0.8836) (Figure 4).

### 3.5. Exploratory atlas-based correlation analysis

This analysis was run post-hoc to gauge the significance of our amygdala finding against other brain regions. In the broad range of regions examined, correlation values ranged from r = 0.46 to r = −0.29 for BAT improvement, from 0.42 to −0.27 for long-term FSQ improvement and from 0.53 to −0.35 for short-term FSQ improvement. For all the clinical variables, the BLA and the whole amygdala are within the most correlated regions out of 155. They are notably the two most correlated regions with BAT improvement as well as the first and fourth with long-term FSQ improvement. The results for all other regions can be found in the supplemental file, Table S1.

## 4. Discussion

Our study aimed to better understand the relationship between the evolution of amygdala activity during exposure and the effectiveness of an exposure intervention. Both an increase and decrease in amygdala activity are compatible with existing theoretical underpinnings from the clinical literature. The habituation model suggests that effective exposure therapy corresponds with reduced fear responses over time, often indicated by decreased amygdala activity. However, one can also interpret such a decrease as a potential sign of cognitive disassociation or lack of engagement in stimuli processing. This perspective aligns with the inhibitory learning model, which emphasizes that forming new associations is more central for successful therapy than observing a reduction of fear responses during the session. The process of establishing these new connections demands an active engagement with the stimuli. From this perspective, a decline in amygdala activity could be viewed unfavorably, signaling a potential disengagement from the therapeutic process. Our data supports that the higher the activity increase, the higher the therapeutic gain. Importantly, the individuals who, during the session, displayed decreasing whole amygdala/BLA activity, i.e., amygdala “habituation”, benefited less from exposure, giving more support for the inhibitory learning model.

We also examined the sub-divisions of the amygdala, specifically the basolateral (BLA) and the central nucleus (CeA) due to their distinct functionalities. First, the BLA mirrored the whole amygdala, which was expected due to the BLA covering most of the whole structure. However, though we expected a decrease in CeA activity to be linked with better clinical outcome, we found no correlation, neither positive nor negative, which may indicate that attenuation of autonomic arousal may not be important for therapy. These findings challenge the concept of amygdala desensitization as a representative marker for clinical improvement, as it would be expected according to the habituation model. Our exploratory analysis of more than 150 other brain regions from the AAL atlas, which revealed the amygdala/BLA as the most correlated regions with one-month FSQ/BAT improvement, also provides insights into the relative importance of various brain regions for exposure, and the salient role of the amygdala/BLA amidst them.

It is worth noticing that we were unable to reproduce the findings of Björkstrand et al. (2020) who reported opposite results, i.e., a positive association between decreased amygdala activity during an exposure session and improvement in spider phobia symptoms. While the fMRI paradigm and the clinical outcome we examined were comparable, significant analysis differences should be noted. Björkstrand et al. (2020) used a voxel-based approach restricted to the amygdala (i.e., small volume correction) to find local clusters that negatively correlated with the clinical outcome. Here, we used a complementary approach, namely averaged activity within pre-defined anatomical regions. The strength of Björkstrand’s et al. voxel-based approach is that it can capture sub-regions of the heterogeneous amygdala nuclei which may not match pre-defined anatomical regions (Kedo et al., 2017). The advantage of using pre- defined anatomical regions is that it is computationally easy to apply to the whole brain and easy to interpret. Another difference was that Björkstrand et al. (2020) seem to have primarily examined the link between activity decrease and clinical improvement without mentioning a potential examination of activity increases, which might have revealed other amygdala clusters. For our analysis, we took into account activity de- and increases.

In addition to informing theoretical accounts of exposure therapy through neuroscientific observations, our study has practical implications for the development of novel brain-based therapeutic approaches. For example, real-time neurofeedback procedures often rely on fMRI-based markers of a psychiatric condition which were defined by contrasting the behavior of a “normal” population with a “clinical” population. For instance, overactivity in the insula has been identified as characteristic of OCD, leading some studies to train patients to downregulate this region (Buyukturkoglu et al., 2015). Similarly, reduced amygdala reactivity to positive stimuli in depression patients has been a target for training (Young et al., 2017). While some of these approaches have yielded clinical improvements, our study suggests caution: guiding a patient’s brain activity towards a “normal” pattern may not always be beneficial. If one uses this logic to design a neurofeedback-based exposure therapy, one would train patients to downregulate their “abnormally” high responses to spider images. Our study, though not causally conclusive, suggests that down-regulation of the amygdala might rather be counterproductive. These insights underscore the critical importance of establishing models of how therapeutic interventions work.

### 4.1. Limitations

One limitation is the fMRI scanning resolution in general: unlike in rodents, the CeA of humans is a very small structure. Smaller regions are more difficult to align in normalized space and it is reasonable to assume that our CeA mask may not have perfectly covered the actual CeA region of each participant. Another limitation is the fact that our design included only one, relatively short, session. Although some therapeutic protocols include only one session as well (Öst, 1989), subsequent amygdala trajectories may look different had there been more runs or more than one session. Also, our cue-exposure task cannot be equated to actual exposure therapy, and participants were not asked to perform any kind of mental task other than to pay attention to the stimulus. Finally, our sample consisted of spider-fearful individuals but not diagnosed phobia patients seeking therapy, and we do not know whether the amygdala of individuals with diagnosed phobia and motivation to improve their symptoms would behave differently.

## 5. Conclusion

To conclude, our study challenges the traditional view that reduced amygdala activity is a hallmark of effective exposure therapy. Instead, our findings suggest that maintained or increased amygdala activity might be crucial for therapeutic success, emphasizing the importance of active engagement with the stimuli. Importantly, the evolution of the amygdala during exposure was one of the most predictive of clinical improvement as compared to more than 150 other predefined atlas regions which further highlights the importance of the amygdala in fear attenuation processes. Our findings also serve as a reminder of the complexities involved when translating neuroscientific insights into brain-based interventions. Caution is advised when leveraging neural markers for direct clinical applications without a comprehensive understanding of their causal relationships and therapeutic implications. Advancing theoretical accounts of interventions, together with a more comprehensive understanding of how they work, is crucial to improving existing therapies and guiding the development of future interventions.

## Acknowledgments

This work was supported by the Swiss National Science Foundation (BSSG10_155915, 100014_178841, 32003B_166566), and the Foundation for Research in Science and the Humanities at the University of Zurich (STWF-17-012). Kathrin Kostorz is supported by the Austrian Science Fund (FWF) ESPRIT Programme (ESP 289). We thank Sophia Shea, Christoph Frühlinger, Marie-Therese Mang, David Gomola, Lisa Neuner, and Elif Dinc for their help in data collection, and Christoph Frühlinger for sketching figures.

ChatGPT and DeepL were used for language improvement and condensation of our original text. The authors reviewed and edited the content as needed and take full responsibility for the content of the publication.

