## Supplemental file for "Amygdala habituation during exposure is associated with failure to reduce phobic symptoms"

Table S1 – Correlation between change of activity in brain regions of the AAL atlas and improvement of spider fear symptoms (Behavioral Avoidance Test one month after exposure (BAT), Fear of Spider Questionnaire (long-term FSQ) after one month, and FSQ after one week (short-term FSQ)).

| Brain region | Correlation with BAT (r) | Significance (p) |
|---|---|---|
| **Amygdala-whole** | **0.458351** | **0.001948998** |
| **Amygdala-BLA** | **0.4508809** | **0.002345928** |
| Thal_Re_R | 0.4250881 | 0.00432318 |
| Thal_LGN_R | 0.4215645 | 0.004684056 |
| Amygdala_R | 0.3787174 | 0.011691686 |
| Hippocampus_R | 0.3761804 | 0.012301153 |
| Thal_PuI_R | 0.3715292 | 0.013489634 |
| Occipital_Sup_L | 0.3465821 | 0.021682874 |
| ParaHippocampal_L | 0.3226216 | 0.0331732 |
| Thal_IL_R | 0.3219168 | 0.033575979 |
| Thal_MGN_R | 0.3219168 | 0.033575979 |
| Amygdala_L | 0.3148696 | 0.037831835 |
| Thal_VPL_R | 0.3033122 | 0.045770591 |
| Temporal_Pole_Mid_R | 0.2796335 | 0.066300847 |
| OFCmed_R | 0.2778013 | 0.068156013 |
| Thal_VA_L | 0.2644116 | 0.083003723 |
| SN_pc_R | 0.2588931 | 0.097874175 |
| Olfactory_R | 0.2546864 | 0.095302643 |
| Thal_Re_L | 0.2535588 | 0.096815857 |
| ParaHippocampal_R | 0.2517266 | 0.099314596 |
| Rectus_L | 0.2466526 | 0.106495574 |
| Hippocampus_L | 0.2408739 | 0.115154112 |
| VTA_L | 0.2386354 | 0.127844899 |
| Lingual_L | 0.2360817 | 0.122734357 |
| Occipital_Sup_R | 0.2339676 | 0.126196648 |
| Red_N_R | 0.2279394 | 0.146234015 |
| Frontal_Med_Orb_R | 0.2264975 | 0.139023943 |
| Vermis_9 | 0.2188865 | 0.153072926 |
| Cuneus_L | 0.2121212 | 0.166419215 |
| N_Acc_L | 0.2116984 | 0.167280744 |
| Calcarine_L | 0.207611 | 0.175777431 |
| Cuneus_R | 0.2040874 | 0.18334982 |

| | | |
|---|---|---|
| SN_pc_L | 0.2020096 | 0.198850729 |
| Temporal_Pole_Sup_R | 0.1985906 | 0.195627608 |
| Thal_MDl_L | 0.1937984 | 0.206800338 |
| Vermis_8 | 0.1936575 | 0.207135628 |
| Thal_MDm_L | 0.1880197 | 0.220863252 |
| Lingual_R | 0.1843552 | 0.230119339 |
| Thal_VPL_L | 0.1811135 | 0.238528196 |
| SN_pr_L | 0.1791589 | 0.255316214 |
| Fusiform_R | 0.1764623 | 0.250957704 |
| Olfactory_L | 0.1737844 | 0.258310131 |
| Calcarine_R | 0.1698379 | 0.269407725 |
| LC_L | 0.1691332 | 0.271422432 |
| Thal_PuL_R | 0.161945 | 0.292545409 |
| Putamen_L | 0.1601128 | 0.298096918 |
| Occipital_Mid_L | 0.1598309 | 0.298957033 |
| Paracentral_Lobule_R | 0.1575758 | 0.305895914 |
| ACC_sub_L | 0.1530655 | 0.320082774 |
| Precuneus_L | 0.1502467 | 0.329158683 |
| Thal_LGN_L | 0.1479915 | 0.336535049 |
| OFCpost_R | 0.1472868 | 0.338861214 |
| Putamen_R | 0.1444679 | 0.348265945 |
| Vermis_6 | 0.1444679 | 0.348265945 |
| LC_R | 0.1424947 | 0.354944337 |
| Temporal_Mid_R | 0.1424947 | 0.354944337 |
| Vermis_4_5 | 0.1420719 | 0.356385587 |
| Fusiform_L | 0.1379845 | 0.370502085 |
| Thal_VL_L | 0.1371388 | 0.373464355 |
| Thal_PuL_L | 0.1361522 | 0.376938309 |
| Raphe_M | 0.1351656 | 0.380431582 |
| Precentral_R | 0.134179 | 0.383944138 |
| Thal_IL_L | 0.134179 | 0.383944138 |
| Raphe_D | 0.1340381 | 0.384447503 |
| SN_pr_R | 0.1325883 | 0.395387778 |
| OFCpost_L | 0.1315011 | 0.3935751 |
| VTA_R | 0.1307762 | 0.40192436 |
| Precuneus_R | 0.1279774 | 0.406461955 |
| Frontal_Inf_Tri_R | 0.1268499 | 0.410636923 |
| Thal_PuI_L | 0.1267089 | 0.411160531 |
| Temporal_Inf_L | 0.1262861 | 0.412733667 |
| Frontal_Med_Orb_L | 0.1238901 | 0.421713431 |
| Thal_PuM_R | 0.1224806 | 0.427047299 |
| **Amygdala-CeN** | **0.1186751** | **0.441638283** |
| Temporal_Pole_Mid_L | 0.1154334 | 0.45428365 |

| | | |
|---|---|---|
| ACC_sub_R | 0.1150106 | 0.455947558 |
| Cingulate_Post_L | 0.1107822 | 0.472768842 |
| Frontal_Inf_Orb_2_R | 0.1071177 | 0.487611946 |
| Temporal_Mid_L | 0.1069767 | 0.48818768 |
| Pallidum_L | 0.1006342 | 0.514460766 |
| OFCant_L | 0.093587 | 0.544469302 |
| Pallidum_R | 0.091191 | 0.554861505 |
| Thal_MGN_L | 0.0882311 | 0.567827753 |
| Rectus_R | 0.085976 | 0.577800539 |
| Vermis_10 | 0.085976 | 0.577800539 |
| Thal_PuA_R | 0.0831572 | 0.590378098 |
| Caudate_R | 0.0823115 | 0.594175141 |
| Temporal_Sup_R | 0.078365 | 0.612036538 |
| Paracentral_Lobule_L | 0.0782241 | 0.612678701 |
| Frontal_Inf_Oper_L | 0.070895 | 0.646460782 |
| Cingulate_Mid_L | 0.0694856 | 0.653042289 |
| Vermis_7 | 0.0692037 | 0.65436177 |
| Cingulate_Post_R | 0.0673714 | 0.662963883 |
| OFClat_L | 0.0645525 | 0.676282459 |
| Occipital_Mid_R | 0.0628612 | 0.68432151 |
| Temporal_Inf_R | 0.0624383 | 0.686336768 |
| Precentral_L | 0.0591966 | 0.701858338 |
| Red_N_L | 0.0585926 | 0.708171624 |
| Insula_R | 0.0553911 | 0.720234574 |
| Thal_MDm_R | 0.0544045 | 0.725025073 |
| Thal_VL_R | 0.0536998 | 0.728453295 |
| ACC_pre_L | 0.053277 | 0.730512774 |
| Thal_PuM_L | 0.053277 | 0.730512774 |
| Frontal_Inf_Oper_R | 0.0525722 | 0.733949438 |
| Thal_PuA_L | 0.0435518 | 0.778375731 |
| Supp_Motor_Area_R | 0.0418605 | 0.786789597 |
| Parietal_Sup_L | 0.040592 | 0.793115994 |
| Frontal_Sup_Medial_L | 0.0341085 | 0.82564994 |
| ACC_pre_R | 0.0336857 | 0.827782514 |
| Thal_LP_L | 0.0312896 | 0.839890021 |
| Caudate_L | 0.0273432 | 0.859911141 |
| Temporal_Pole_Sup_L | 0.012544 | 0.93567245 |
| Supp_Motor_Area_L | 0.0108527 | 0.944379983 |
| Parietal_Sup_R | 0.0105708 | 0.945831895 |
| N_Acc_R | 0.0093023 | 0.9523676 |
| Frontal_Inf_Orb_2_L | 0.0060606 | 0.969083124 |
| Cingulate_Mid_R | -0.001832 | 0.990904777 |
| Thal_AV_L | -0.003805 | 0.980719547 |

| | | |
|---|---|---|
| Frontal_Sup_2_L | -0.004228 | 0.978537341 |
| Frontal_Sup_Medial_R | -0.006765 | 0.965447942 |
| OFCant_R | -0.00747 | 0.961813459 |
| OFCmed_L | -0.012403 | 0.936397805 |
| Occipital_Inf_L | -0.013108 | 0.932771552 |
| Cingulate_Ant_L | -0.013249 | 0.932046462 |
| Cingulate_Ant_R | -0.013249 | 0.932046462 |
| Thalamus_L | -0.013249 | 0.932046462 |
| Thalamus_R | -0.013249 | 0.932046462 |
| Vermis_3 | -0.020014 | 0.897317392 |
| Rolandic_Oper_R | -0.020296 | 0.895874039 |
| Frontal_Inf_Tri_L | -0.020719 | 0.893709641 |
| ACC_sup_L | -0.028048 | 0.856329127 |
| SupraMarginal_R | -0.040592 | 0.793115994 |
| Postcentral_R | -0.047921 | 0.756758805 |
| Postcentral_L | -0.06568 | 0.670942908 |
| Thal_MDl_R | -0.066244 | 0.668279164 |
| Temporal_Sup_L | -0.072023 | 0.641214864 |
| Rolandic_Oper_L | -0.073714 | 0.633378641 |
| ACC_sup_R | -0.08513 | 0.581560901 |
| OFClat_R | -0.085271 | 0.580933401 |
| SupraMarginal_L | -0.093164 | 0.546296359 |
| Vermis_1_2 | -0.093305 | 0.545687011 |
| Heschl_R | -0.101339 | 0.511506636 |
| Insula_L | -0.112474 | 0.466000745 |
| Occipital_Inf_R | -0.113319 | 0.462636439 |
| Frontal_Mid_2_L | -0.122058 | 0.428654887 |
| Thal_AV_R | -0.127273 | 0.409068413 |
| Parietal_Inf_L | -0.129105 | 0.402311747 |
| Thal_VA_R | -0.131642 | 0.393064686 |
| Frontal_Sup_2_R | -0.134038 | 0.384447503 |
| Thal_LP_R | -0.14024 | 0.362672384 |
| Angular_R | -0.148555 | 0.334681331 |
| Angular_L | -0.166455 | 0.279169706 |
| Frontal_Mid_2_R | -0.175617 | 0.253264005 |
| Heschl_L | -0.195913 | 0.201816902 |
| Parietal_Inf_R | -0.291896 | 0.054902688 |

| Brain region | Correlation with long-term FSQ (r) | Significance (p) |
|---|---|---|
| **Amygdala-BLA** | **0.417191341** | **0.0043565** |
| Red_N_R | 0.392671039 | 0.00919857 |
| Frontal_Med_Orb_R | 0.387924007 | 0.00846244 |

| | | |
|---|---|---|
| **Amygdala-whole** | **0.376783964** | **0.01073495** |
| Cuneus_L | 0.370983231 | 0.01211276 |
| VTA_L | 0.361692566 | 0.01716303 |
| Amygdala_R | 0.359315848 | 0.0153462 |
| Hippocampus_R | 0.349955574 | 0.01844464 |
| Red_N_L | 0.332922045 | 0.02723056 |
| Cingulate_Post_R | 0.293068842 | 0.05072728 |
| SN_pc_R | 0.289006489 | 0.06015867 |
| Amygdala_L | 0.278698845 | 0.06375641 |
| ParaHippocampal_L | 0.278501093 | 0.06395272 |
| ParaHippocampal_R | 0.271777516 | 0.07091865 |
| Rectus_L | 0.259253207 | 0.0854825 |
| Frontal_Med_Orb_L | 0.25226596 | 0.09456547 |
| Thal_PuI_L | 0.243301191 | 0.10729082 |
| Occipital_Sup_L | 0.239082477 | 0.11371404 |
| Thal_LGN_R | 0.237830046 | 0.11567606 |
| Temporal_Pole_Mid_R | 0.237236789 | 0.11661435 |
| OFCpost_R | 0.230710964 | 0.1273195 |
| VTA_R | 0.223052834 | 0.14556453 |
| Olfactory_R | 0.221021104 | 0.14455245 |
| Hippocampus_L | 0.218911747 | 0.14852215 |
| Temporal_Pole_Mid_L | 0.214099775 | 0.15787758 |
| Occipital_Sup_R | 0.2104084 | 0.16534091 |
| Cuneus_R | 0.20605785 | 0.17446187 |
| Temporal_Mid_L | 0.202102805 | 0.1830632 |
| ACC_pre_L | 0.19992753 | 0.18792112 |
| SN_pc_L | 0.19796 | 0.20319699 |
| Cingulate_Post_L | 0.196697577 | 0.19530257 |
| Frontal_Sup_Medial_L | 0.196499824 | 0.19576107 |
| Olfactory_L | 0.188655652 | 0.2145656 |
| LC_L | 0.176724599 | 0.24551479 |
| OFCpost_L | 0.176526847 | 0.24605195 |
| Raphe_D | 0.176395012 | 0.2464105 |
| OFCmed_R | 0.170857949 | 0.26178896 |
| ACC_sub_L | 0.167693912 | 0.27085787 |
| Frontal_Sup_Medial_R | 0.167364325 | 0.27181435 |
| N_Acc_L | 0.157476713 | 0.30154646 |
| Lingual_L | 0.152335154 | 0.31780205 |
| Temporal_Pole_Sup_R | 0.152203319 | 0.31822601 |
| Frontal_Inf_Orb_2_L | 0.144820568 | 0.34253692 |
| Occipital_Mid_R | 0.141326945 | 0.3544298 |
| Lingual_R | 0.136449056 | 0.37144984 |
| Paracentral_Lobule_R | 0.133944194 | 0.38037664 |

| | | |
|---|---|---|
| Raphe_M | 0.132098507 | 0.38703489 |
| Angular_L | 0.130516488 | 0.39279616 |
| OFCmed_L | 0.128604883 | 0.39982417 |
| Rectus_R | 0.128275296 | 0.40104322 |
| LC_R | 0.126100021 | 0.40914281 |
| Frontal_Sup_2_L | 0.12484759 | 0.41384848 |
| Temporal_Pole_Sup_L | 0.123265572 | 0.41983641 |
| OFClat_L | 0.117992179 | 0.44014726 |
| SN_pr_L | 0.116207054 | 0.45803539 |
| Thal_PuI_R | 0.108895575 | 0.47642806 |
| Paracentral_Lobule_L | 0.104281356 | 0.49541748 |
| Occipital_Mid_L | 0.102699338 | 0.50201669 |
| Thal_PuM_L | 0.101908329 | 0.50533303 |
| Thal_MDl_R | 0.101249155 | 0.50810513 |
| Calcarine_R | 0.091954799 | 0.54799432 |
| Fusiform_R | 0.091625212 | 0.54943574 |
| ACC_pre_R | 0.090109111 | 0.5560894 |
| Thal_MDm_R | 0.088988515 | 0.56103156 |
| SN_pr_R | 0.080180548 | 0.60489052 |
| Vermis_9 | 0.079825994 | 0.60219028 |
| Thal_PuM_R | 0.076727875 | 0.61639795 |
| Temporal_Inf_R | 0.07659604 | 0.61700567 |
| Thal_VPL_L | 0.074025261 | 0.62890634 |
| Thal_MGN_L | 0.070795307 | 0.64399102 |
| Thal_MGN_R | 0.070267968 | 0.64646752 |
| ACC_sup_L | 0.069938381 | 0.64801726 |
| Precuneus_R | 0.069476959 | 0.65018936 |
| Angular_R | 0.06894962 | 0.65267528 |
| Calcarine_L | 0.068356363 | 0.6554764 |
| Thal_MDm_L | 0.064401318 | 0.67426855 |
| Temporal_Mid_R | 0.060709942 | 0.69198683 |
| Thal_LP_R | 0.056820815 | 0.7108317 |
| OFCant_L | 0.056491228 | 0.71243678 |
| Fusiform_L | 0.054777375 | 0.72080286 |
| Occipital_Inf_R | 0.053129439 | 0.72887769 |
| Thal_LP_L | 0.051877008 | 0.73503409 |
| N_Acc_R | 0.050229073 | 0.74315964 |
| Thal_PuL_R | 0.046076275 | 0.76375699 |
| ACC_sub_R | 0.033486049 | 0.82713988 |
| ACC_sup_R | 0.03269504 | 0.83116312 |
| Frontal_Inf_Orb_2_R | 0.025905545 | 0.86586135 |
| Thal_IL_R | 0.02017073 | 0.8953679 |
| Occipital_Inf_L | 0.019445638 | 0.89910942 |

| | | |
|---|---|---|
| Thal_MDl_L | 0.01430408 | 0.92569765 |
| Vermis_4_5 | 0.009096604 | 0.95270848 |
| Thal_Re_R | 0.00408688 | 0.97874349 |
| Temporal_Inf_L | 0.002504862 | 0.98697089 |
| Precuneus_L | 0.000197752 | 0.99897134 |
| Thal_IL_L | 6.59E-05 | 0.99965711 |
| Thal_VPL_R | -0.001845688 | 0.99039942 |
| Thal_Re_L | -0.003955045 | 0.97942904 |
| OFCant_R | -0.005471146 | 0.97154628 |
| Thal_PuA_R | -0.005800733 | 0.96983297 |
| Vermis_7 | -0.00995353 | 0.94825925 |
| Postcentral_R | -0.011074126 | 0.94244339 |
| Thal_LGN_L | -0.015688346 | 0.91853019 |
| Frontal_Sup_2_R | -0.018918299 | 0.90183189 |
| Thal_AV_R | -0.021554996 | 0.8882313 |
| Thal_VA_R | -0.025773711 | 0.86653779 |
| Caudate_R | -0.028674077 | 0.85167859 |
| Parietal_Sup_L | -0.028805912 | 0.85100434 |
| Precentral_R | -0.029465086 | 0.84763466 |
| Thal_VL_L | -0.032629122 | 0.83149859 |
| Thal_PuL_L | -0.033749718 | 0.82579976 |
| Parietal_Sup_R | -0.043901001 | 0.77461181 |
| OFClat_R | -0.044362423 | 0.77230564 |
| Thal_VL_R | -0.046537697 | 0.76146012 |
| Postcentral_L | -0.047130954 | 0.75850998 |
| Caudate_L | -0.051151917 | 0.73860587 |
| **Amygdala-CeN** | **-0.052931687** | **0.72984865** |
| Cingulate_Mid_L | -0.054711457 | 0.72112528 |
| Frontal_Inf_Tri_R | -0.062489713 | 0.68342307 |
| Vermis_8 | -0.062687465 | 0.68247394 |
| Frontal_Inf_Oper_L | -0.066181088 | 0.66578696 |
| Thal_VA_L | -0.06690618 | 0.66234321 |
| Frontal_Mid_2_L | -0.066972097 | 0.66203049 |
| Thal_PuA_L | -0.069806546 | 0.64863757 |
| Frontal_Inf_Tri_L | -0.074025261 | 0.62890634 |
| Heschl_R | -0.075805031 | 0.62065728 |
| Parietal_Inf_L | -0.075936866 | 0.62004805 |
| Temporal_Sup_L | -0.081144342 | 0.59618797 |
| Precentral_L | -0.09472333 | 0.53595809 |
| Cingulate_Mid_R | -0.108961493 | 0.4761596 |
| Vermis_1_2 | -0.109159245 | 0.4753547 |
| Vermis_6 | -0.120035619 | 0.43221306 |
| Putamen_R | -0.120167454 | 0.43170394 |

| | | |
|---|---|---|
| Heschl_L | -0.128077544 | 0.40177569 |
| Cingulate_Ant_L | -0.13302135 | 0.38369724 |
| Cingulate_Ant_R | -0.13302135 | 0.38369724 |
| Thalamus_L | -0.13302135 | 0.38369724 |
| Thalamus_R | -0.13302135 | 0.38369724 |
| Temporal_Sup_R | -0.136580891 | 0.37098351 |
| Insula_R | -0.137899239 | 0.3663395 |
| Frontal_Inf_Oper_R | -0.144293229 | 0.34431612 |
| Vermis_3 | -0.145875247 | 0.33899558 |
| Vermis_10 | -0.148511944 | 0.33024179 |
| Rolandic_Oper_L | -0.149830292 | 0.32591833 |
| Rolandic_Oper_R | -0.151346393 | 0.32099045 |
| Putamen_L | -0.156685704 | 0.30401192 |
| Insula_L | -0.174219737 | 0.25237752 |
| Pallidum_R | -0.175142581 | 0.2498343 |
| Frontal_Mid_2_R | -0.176197259 | 0.24694898 |
| Supp_Motor_Area_R | -0.183645928 | 0.22721267 |
| Supp_Motor_Area_L | -0.186809964 | 0.2191668 |
| SupraMarginal_L | -0.188919321 | 0.21391381 |
| Parietal_Inf_R | -0.198609182 | 0.19090959 |
| SupraMarginal_R | -0.250354355 | 0.09717597 |
| Pallidum_L | -0.260835225 | 0.0835237 |
| Thal_AV_L | -0.268745315 | 0.07425032 |

| Brain region | Correlation with short-term FSQ (r) | Significance (p) |
|---|---|---|
| Occipital_Sup_L | 0.5336182 | 0.00056054 |
| Occipital_Sup_R | 0.50219025 | 0.00131504 |
| Occipital_Mid_R | 0.43911533 | 0.00581367 |
| Calcarine_R | 0.43550166 | 0.00627929 |
| Occipital_Mid_L | 0.43265453 | 0.00666845 |
| Hippocampus_R | 0.42597472 | 0.00766378 |
| Fusiform_R | 0.42258006 | 0.00821669 |
| Cuneus_L | 0.41798085 | 0.00902006 |
| Lingual_R | 0.40768737 | 0.01106443 |
| Occipital_Inf_L | 0.39005706 | 0.01548163 |
| ParaHippocampal_R | 0.38852399 | 0.01592761 |
| Occipital_Inf_R | 0.37757348 | 0.01943837 |
| **Amygdala-whole** | **0.37494536** | **0.02037101** |
| Cuneus_R | 0.37483585 | 0.02041067 |
| Fusiform_L | 0.36826555 | 0.02291092 |
| Lingual_L | 0.36344733 | 0.02490206 |
| **Amygdala-BLA** | **0.34625503** | **0.03321081** |
| Rectus_L | 0.32369699 | 0.04742533 |

| | | |
|---|---|---|
| Thal_LGN_R | 0.31822173 | 0.05152627 |
| Amygdala_R | 0.31767421 | 0.05195153 |
| Calcarine_L | 0.29697775 | 0.07019757 |
| Amygdala_L | 0.28482269 | 0.08305718 |
| Red_N_L | 0.26700676 | 0.11014291 |
| Vermis_7 | 0.26445474 | 0.10861196 |
| ParaHippocampal_L | 0.25755592 | 0.11849709 |
| Thal_PuI_R | 0.25186166 | 0.12715297 |
| Hippocampus_L | 0.23039867 | 0.16403356 |
| Frontal_Med_Orb_R | 0.21616301 | 0.19241116 |
| Olfactory_L | 0.21320637 | 0.19871356 |
| Thal_IL_L | 0.21254934 | 0.20013349 |
| Red_N_R | 0.21223443 | 0.21398533 |
| Temporal_Inf_R | 0.20882617 | 0.20831381 |
| Caudate_R | 0.20006576 | 0.22846908 |
| Parietal_Sup_R | 0.18418753 | 0.26830097 |
| Vermis_4_5 | 0.18407802 | 0.26859061 |
| Rectus_R | 0.17739821 | 0.28664736 |
| Olfactory_R | 0.17378455 | 0.29673538 |
| OFClat_L | 0.16743325 | 0.31500995 |
| Frontal_Med_Orb_L | 0.16272453 | 0.32900551 |
| Thal_MDl_R | 0.16042493 | 0.33597858 |
| Thal_MDm_L | 0.16020592 | 0.3366474 |
| Temporal_Pole_Sup_L | 0.15965839 | 0.33832303 |
| Caudate_L | 0.15045997 | 0.36723629 |
| OFCpost_L | 0.14662729 | 0.37970563 |
| Thal_LP_R | 0.14169956 | 0.39609865 |
| Thal_PuI_L | 0.13797639 | 0.40875136 |
| VTA_L | 0.13689636 | 0.4259454 |
| Raphe_D | 0.1360053 | 0.41554202 |
| Frontal_Inf_Orb_2_L | 0.13217262 | 0.4289271 |
| Temporal_Pole_Sup_R | 0.12231716 | 0.46442521 |
| Thal_MDm_R | 0.12012706 | 0.47252067 |
| Thal_MGN_R | 0.11804646 | 0.48027973 |
| Temporal_Pole_Mid_L | 0.11465181 | 0.49308063 |
| OFCmed_R | 0.11147616 | 0.50521215 |
| Temporal_Pole_Mid_R | 0.10698645 | 0.52261684 |
| N_Acc_L | 0.09636446 | 0.56493366 |
| OFCmed_L | 0.09592644 | 0.56671196 |
| ACC_sub_L | 0.09242228 | 0.58103001 |
| Paracentral_Lobule_R | 0.08716604 | 0.60280551 |
| LC_L | 0.08519494 | 0.61106098 |
| Thal_PuL_R | 0.08344286 | 0.61843918 |

| | | |
|---|---|---|
| SN_pr_L | 0.08074696 | 0.63967785 |
| OFCant_L | 0.0798292 | 0.63377296 |
| Vermis_9 | 0.07873414 | 0.63844984 |
| Thal_PuM_R | 0.07687256 | 0.64643217 |
| SN_pc_L | 0.07289119 | 0.6726782 |
| Temporal_Inf_L | 0.07183533 | 0.66822555 |
| LC_R | 0.07161632 | 0.66917935 |
| Putamen_R | 0.07095928 | 0.67204383 |
| Thal_PuA_R | 0.06548403 | 0.69608868 |
| SN_pc_R | 0.06117194 | 0.72303117 |
| Thal_LGN_L | 0.06066581 | 0.71749385 |
| Parietal_Sup_L | 0.0583662 | 0.72778653 |
| Thal_VA_R | 0.05584759 | 0.73911359 |
| Raphe_M | 0.04894877 | 0.77041222 |
| Thal_IL_R | 0.04774421 | 0.77591554 |
| Vermis_6 | 0.04675866 | 0.7804263 |
| ACC_pre_L | 0.04007886 | 0.81117895 |
| Thal_Re_L | 0.03690321 | 0.82590057 |
| OFCpost_R | 0.03383707 | 0.84017044 |
| Thal_LP_L | 0.03197548 | 0.84885921 |
| Frontal_Sup_Medial_L | 0.03131845 | 0.85193009 |
| Thal_VL_R | 0.02562419 | 0.87862981 |
| ACC_sub_R | 0.02124398 | 0.89925998 |
| Putamen_L | 0.01839685 | 0.91270549 |
| Thal_VPL_R | 0.0171923 | 0.91840141 |
| Vermis_8 | 0.01708279 | 0.91891942 |
| Thal_VPL_L | 0.0152212 | 0.92773053 |
| Frontal_Sup_2_L | 0.0151117 | 0.9282491 |
| Temporal_Mid_L | 0.01171704 | 0.94433836 |
| Thal_VL_L | 0.0107315 | 0.94901379 |
| Thal_MGN_L | 0.00733684 | 0.96512997 |
| Thal_PuM_L | 0.00394218 | 0.9812598 |
| Paracentral_Lobule_L | 0.00262812 | 0.98750592 |
| Vermis_3 | 0.00153307 | 0.9927116 |
| Pallidum_R | -0.0013141 | 0.99375278 |
| Thal_VA_L | -0.0045992 | 0.97813713 |
| Thal_MDl_L | -0.0077749 | 0.96304956 |
| Precentral_R | -0.0079939 | 0.96200945 |
| Cingulate_Post_R | -0.0087604 | 0.95836955 |
| Precuneus_R | -0.0116075 | 0.94485777 |
| Cingulate_Post_L | -0.0170828 | 0.91891942 |
| Cingulate_Ant_L | -0.0175208 | 0.91684757 |
| Cingulate_Ant_R | -0.0175208 | 0.91684757 |

| | | |
|---|---|---|
| Thalamus_L | -0.0175208 | 0.91684757 |
| Thalamus_R | -0.0175208 | 0.91684757 |
| Thal_AV_R | -0.0185064 | 0.91218789 |
| Pallidum_L | -0.0239816 | 0.88635761 |
| **Amygdala-CeN** | **-0.0267192** | **0.87348404** |
| VTA_R | -0.0284681 | 0.86716957 |
| SN_pr_R | -0.0291798 | 0.86387943 |
| Postcentral_R | -0.0346036 | 0.83659807 |
| ACC_sup_L | -0.042488 | 0.80005304 |
| Temporal_Sup_L | -0.0549715 | 0.74306632 |
| Precuneus_L | -0.0593517 | 0.7233695 |
| Insula_R | -0.0644985 | 0.70044883 |
| Temporal_Sup_R | -0.0733684 | 0.66156335 |
| Thal_PuL_L | -0.0863995 | 0.60601026 |
| Cingulate_Mid_L | -0.0940649 | 0.57429825 |
| Vermis_1_2 | -0.096693 | 0.56360163 |
| Frontal_Inf_Oper_L | -0.096912 | 0.56271441 |
| Frontal_Inf_Tri_R | -0.1011827 | 0.54554406 |
| Angular_R | -0.1022777 | 0.54118187 |
| OFCant_R | -0.1061104 | 0.52604692 |
| Supp_Motor_Area_L | -0.1120237 | 0.50310983 |
| Thal_Re_R | -0.1131187 | 0.4989185 |
| Heschl_R | -0.1153088 | 0.49058946 |
| N_Acc_R | -0.117937 | 0.48068994 |
| Temporal_Mid_R | -0.129435 | 0.43863296 |
| Thal_AV_L | -0.129654 | 0.43785207 |
| Insula_L | -0.130092 | 0.4362926 |
| Thal_PuA_L | -0.130311 | 0.43551401 |
| Frontal_Inf_Oper_R | -0.1325011 | 0.42777049 |
| ACC_sup_R | -0.1376479 | 0.40987872 |
| Frontal_Inf_Orb_2_R | -0.1399475 | 0.40202441 |
| Postcentral_L | -0.1418091 | 0.39572998 |
| Frontal_Inf_Tri_L | -0.1423566 | 0.39388959 |
| Frontal_Sup_Medial_R | -0.1424661 | 0.39352211 |
| Supp_Motor_Area_R | -0.1442182 | 0.38766948 |
| Angular_L | -0.1483794 | 0.37397474 |
| Rolandic_Oper_L | -0.1529786 | 0.35917688 |
| Rolandic_Oper_R | -0.1574683 | 0.34507671 |
| Cingulate_Mid_R | -0.1708279 | 0.30515624 |
| ACC_pre_R | -0.173456 | 0.29766361 |
| OFClat_R | -0.1870347 | 0.2608426 |
| SupraMarginal_R | -0.1939335 | 0.24334456 |
| Precentral_L | -0.20576 | 0.2152226 |

| | | |
|---|---|---|
| Frontal_Mid_2_L | -0.2121113 | 0.20108405 |
| Frontal_Sup_2_R | -0.2198862 | 0.1846767 |
| Frontal_Mid_2_R | -0.2240474 | 0.17629602 |
| Heschl_L | -0.2359834 | 0.15377006 |
| SupraMarginal_L | -0.2429917 | 0.14156205 |
| Parietal_Inf_L | -0.270149 | 0.10093424 |
| Vermis_10 | -0.2778144 | 0.09126141 |
| Parietal_Inf_R | -0.3486641 | 0.0319254 |
